# Weed communities and wheat yield are modified by cropping systems and climate conditions

**DOI:** 10.1101/2020.07.09.190488

**Authors:** Tim Seipel, Suzanne L. Ishaq, Fabian D. Menalled

## Abstract

Understanding the impact of biological and environmental stresses on crop performance is essential to secure the long-term sustainability of agricultural production. How cropping systems modify weed communities and wheat yield in response to predicted climate conditions is unknown. We tested the effect of warmer, and warmer and drier conditions on weed biomass, weed community characteristics, and winter wheat yields in three contrasting cropping systems: a no-till chemically managed system, a tilled organic system, and an organic system that used grazing to reduce tillage. Weed communities in the organic systems were more diverse and more variable than the no-till conventional system, though the grazed organic and no-till conventional systems had more similar relative species abundance. Cropping system affected weed biomass and weed species composition recorded in 0.75 m^2^ split-plots, with the most biomass recorded in grazed organic system (38 g ±23.4 SE) compared to the tilled-organic (17 g ±10.3 SE) and no-till chemically managed systems (<1 g ±0.02). Climate conditions had relatively minor impacts on weed communities compared with cropping systems. Wheat yield was highest in the no-till conventional system but declined in response to warmer and drier conditions despite its low weed biomass. Yield was lower in the tilled organic and grazed organic cropping system but declines in warmer and drier conditions were more variable among years. In the Northern Great Plains, predicted climate scenarios have the potential to alter weed communities and reduce wheat yield, and designing resilient cropping systems is essential to mitigate these negative impacts.

## Introduction

Agricultural production faces numerous environmental, social, and economic challenges that test its long-term sustainability, especially in the face of unprecedented market and climate conditions. Global demand for food is expected to increase, consumer preferences are shifting, and climate change is impacting agricultural production, potentially reducing global food security (Rosenzweig & Parry, 1994, Scheelbeek et al., 2018, Tigchelaar et al., 2018). For example in the Northern Great Plains of the United States, the region where this study was conducted and a major cereal-producing system, climate change is predicted to increase temperatures by 2.5 – 3.3°C in the next 30 years and decrease precipitation during the summer, resulting in increased summer drought stress (Whitlock et al., 2017). To ensure global food security, it is therefore vital we understand how climate change will impact agricultural production by modifying planting and harvesting dates, crop phenological development, grain volume weight, and overall yield (Lanning et al., 2010, Morgounov et al., 2018). Higher temperatures are expected to reduce agricultural production especially of wheat (*Triticum aestivium* L.), one of the worlds staple commodities (Wilcox & Makowski, 2014). For example, a 1 °C increase in global temperature is estimated to reduce wheat yield by 8% (Tack et al., 2015, Liu et al., 2016).

In addition to direct effects of increasing global temperatures on wheat production, biotic interactions are likely to further decrease yields (Ramesh et al., 2017, Deutsch et al., 2018).Unfavorable climate conditions increases crop stress and decreasing their competitive ability against weeds, further exacerbating yield loss (Tubiello et al., 2007). Climate change could also lead to shifts in the composition of weed communities, potentially altering the intensity of competitive outcomes between crops and weeds (Ramesh et al., 2017). For instance, habitat suitability of *Bromus tectorum* in the Northern Great Plains of the United States, one of the most problematic weeds in winter wheat, is expected to increase under predicted warmer and drier conditions which may lead to greater negative impacts (Stahlman & Miller, 1990, Bradley, 2009).

Different cropping systems have the potential to modify agroecosystem response to stress and disturbance, and impact the efficacy of specific management practices (Menalled et al., 2001, Nkurunziza et al., 2017, Peterson et al., 2018, Ball et al., 2019). Yet, to our knowledge, no study has systematically compared the impact of climate change across different cropping systems. To assess this knowledge gap, we compared how weed species composition, weed biomass, and wheat yield vary in response to climate conditions across three contrasting cropping systems: 1) a no-till chemically managed system, 2) a tilled-organic systems, and 3) a grazed/reduced-till organic system. Our study focuses on two questions 1) Does faming system and climate conditions affect weed biomass and weed communities? and 2) How do farming system and climate conditions affect wheat yields?

## Methods and Materials

### Study Site and Experimental Design

Our study was conducted within fields of a large five-year cropping system experiment that was conducted at the Montana State University’s Fort Ellis Research and Extension Center, located east of Bozeman, Mt, USA (45.6671 latitude, longitude −110.9977, elevation 1500 m a.s.l.). The mean annual temperature at the site is 6.6 °C and mean total annual precipitation is 44 cm (PRISM Climate Group, 2018). Soils at the Fort Ellis site are classified as a silt loam (a fine-silty, mixed, superactive, frigid Typic Arguistoll) with 0 to 4% slopes and consistent ratio of 1 part sand, 2 parts silt, 1 part clay by weight.

The large cropping system experiment had a total of 45 fields (90 × 12 m) equally divided among three cropping systems: 1) a no-till system that relied on synthetic inputs in the form of fertilizers, herbicides, and fungicides to manage nutrient availability, weed abundance, and pathogen pressure, 2) a tilled-organic system reliant on tillage to manage weeds and terminate cover crops, and 3) a grazed/reduced-till organic system that incorporated sheep grazing to manage weeds and terminate cover crops with the overall goal of reducing the need for tillage. Inputs utilized in the non-till system included 2,4D, bromoxynil, dicamba, fluroxypyr, glyphosate, MCPA, pinoxaden, and urea, which are reflective of typical conventional farm management practices in the Northern Great Plains region. Both organic treatments began the organic transition process in July 2012, making crops harvested in 2015 USDA certified as organic. In the organic tilled system, tillage was accomplished using a chisel plow, tandem disk, or field cultivator, as needed for weed control, seedbed preparation, and to incorporate cover crops and crop residues. Weed control was enhanced with a rotary harrow. In the organic grazed system, targeted sheep grazing was used to reduce tillage intensity for pre-seeding and post-harvest weed control and to terminate the cover crops, with duration and intensity of grazing based on weed or cover crop biomass. Within each cropping system, each crop phase of a five-year crop rotation was present each year and replicated three times. The crop rotation sequence was year 1: safflower (*Carthamus tinctorius* L.) undersown with sweet clover (*Melilotus officinalis* L.), year 2: sweet clover cover crop, year 3: winter wheat, year 4: lentil (*Lens culinaris* L.), and year 5 winter wheat. Additional details on the site history and maintenance can be found in (Lehnhoff et al., 2017, Seipel et al., 2019).

This experiment was conducted in the third year of the rotation where winter wheat was grown in the 2015-2016 and 2016-2017 growing seasons. During the experiment average monthly temperature and precipitation during the spring growing period (March through June) in both 2016 and 2017 were slightly above average (0.5 to 1.5 °C), compared with 30-yr mean monthly temperatures from 1981-2010 (PRISM Climate Group, 2018). In 2016 precipitation was slightly below normal during the March through June period, but in 2017 was above normal (Appdendix Table S2).

Within each winter wheat field, we randomly established three 0.75 m^2^ split-plots, and assigned them to one of three climate treatments: 1) ambient temperature and moisture, used as control, 2) warmer treatment that increased air and soil temperature using open-top chambers, and 3) a warmer and drier treatment that increased temperature and decreased rainfall using a combination of rainout shelters and open-top chambers. Open-top chambers followed (Marion et al., 1997) and were constructed of 1 mm thick Sun-Lite HP (Solar Components Corporation) with a basal diameter of 1.6 m and the top opening diameter was 1.0 m. The height of the chambers was 0.5 m, and chamber wall had an incline of 65°. Rainout shelters were used to reduce the amount of moisture that fell onto split plots with by approximately 50% (Yahdjian & Sala, 2002). Rainout shelters were constructed with a wooden frame and corrugated polycarbonate plastic that covered approximately 50% of a 2 m by 2 m area centered over the warmer and drier split-plots (Appendix S1 Fig 1). The rainout shelter was oriented west to east; the west side was lower and faced into the prevalent wind, and the incline of the rainout shelter increased from west to east by approximately 30°. These climate manipulation structures were placed in the field as winter wheat emerged from dormancy in early spring (early March 2016 and early April 2017) and were removed after of harvest of wheat in early August. To monitor the impact climate manipulations we recorded soil moisture using Delmhorst gypsum block sensors (https://www.delmhorst.com; accessed May 1, 2019) and monitored soil temperature 5 cm below the surface using Maxim ibuttons (https://www.maximintegrated.com/; accessed May 1, 2019).

### Weed Biomass and Wheat Yield Sampling

In each split plot, we destructively sampled aboveground weed biomass in mid-July by cutting the weeds as they began to senesce and no longer had a competitive impact on the ripening wheat. Within each 0.75 m^2^ circular split plot, we cut the weed biomass at ground level and separated each species. The individual biomass of each species was dried and weighed. We left the wheat crop in the field and harvested when it had completely senesced and ripened. To harvest winter wheat, which occurred during the first the last week of July in 2016 and the first week in August in 2017, we cut the two center 75 cm rows of the split plot for a total of 1.5 row meters per split plot. The total biomass of wheat was weighed, and grain was threshed and then weighed. We recorded the biomass of each individual weed species and the grain yields in each split-plot.

### Data Analysis

To assess how climate manipulations affected soil temperature and soil moisture across the three cropping systems, we used general additive models to assess differences throughout the growing season, using day as date and time as the predictor (Wood, 2017). Soil moisture was measured as electrical conductivity and then converted to soil water potential expressed in bars based on conversion table in the <u>Delmhorst model KS-D1 owner’s manual</u>.

We fit linear mixed-effects models using ‘lmerTest’ package in R (Kuznetsova et al., 2017, R Core Team, 2017) to assess differences in the number of weed species (richness), Simpson’s diversity metric, and total weed biomass in response to climate condition, cropping system, and their interaction with year where block was fit as a random factor. The data was log transformed to reduce skew, and all results were back transformed. Post-hoc separation of means was conducted using ‘emmeans’ package and figures were produced using the ‘ggplot2’ package (Wickham, 2009, Lenth, 2019). We compared the differences in weed species biomass among split-plots with PERMANOVA using the ‘adonis’ function in the ‘vegan’ package in R (Oksanen et al., 2017). We used the Bray-Curtis dissimilarity measure for weed biomass as the response variables. In the PERMANOVA, we used cropping system, climate conditions, and their interactions as predictor variables. Year was used to constrain variation within trials of the experiment. We used principal coordinates analysis and bi-plots to visualize differences in dissimilarity among predictor variables based on the results of PERMANOVA.

To compare the impact of cropping system and climate conditions on wheat yield, we fit linear mixed-effects regression models of wheat yield in split-plots in response to cropping system, climate treatments, year, and their interactions, with plot as random effect. We assessed the effects of the predictor variables using Type III analysis of variance. If these variables or their interactions explained variation in wheat yield, we conducted post-hoc differences among estimated means using ‘emmeans’.

## Results

### Temperature and moisture manipulations

Open-top chambers and rain-out shelters reduced soil moisture and increased soil temperature, relative to ambient conditions, during the critical spring growing season when wheat tillers, heads, and kernels develop (date x climate condition interaction P=0.015; Appendix Fig 1-3). In 2016, climate manipulations had a greater effect on soil moisture and soil temperature compared to 2017. In 2016 for the period of early March through the end of June average soil temperatures were 8.6, 8.7 and 8.9 °C in ambient, warmer, and warmer and drier split-plots, respectively. In 2017, average early March through the end of June soil temperatures were 7.5, 7.6 and 7.8 °C in ambient, warmer, and warmer and drier conditions. Under warmer and drier conditions soil moisture was two times less than ambient conditions or warmer condition, especially in in 2016 (Appendix Fig 2 and Fig 3). During 2017 soil moisture in the warmer and drier was lower during the early March to late June, but ample moisture resulted in a smaller intra-year difference and higher available moisture compared to 2016.

### Weed community response

Weed species richness varied among cropping systems (X^2^= 10, P=0.007), and was greatest in the grazed organic cropping system which had a total of 16 species and an average of 4.8 species (±1.2 SE) per 0.75 m^2^ split-plot (Fig 1). A total of 10 species were recorded in all tilled organic split plots, and an average of 2.2 species (±1.2 SE) were recorded in each split plot per year. The no-till conventionally managed cropping system had the lowest number of weed species (3 total) and the lowest average of number of weed species 1.1 (±0.2 SE). Species richness did not differ in response of the year of the trial (X^2^ =0.07, 0.80). Inverse Simpson’s diversity index also varied among cropping systems (F=32, P= <0.001), and was also greatest in the grazed organic cropping system, but the tilled organic and no-till conventional cropping systems did not differ (t= - 1.816, P=0.17; Fig. 1). Diversity did not vary between years (F=2.4, P=0.12). Manipulated climate conditions did not affect weed species richness (X^2^=0.12, P 0.94), Simpson’s diversity measure (F=1.14, P=0.32), or interact with cropping systems (richness X^2^= 3.9621, P=0.41; and Simpson’s diversity F=1.11, P=0.36).

**Fig 1.**
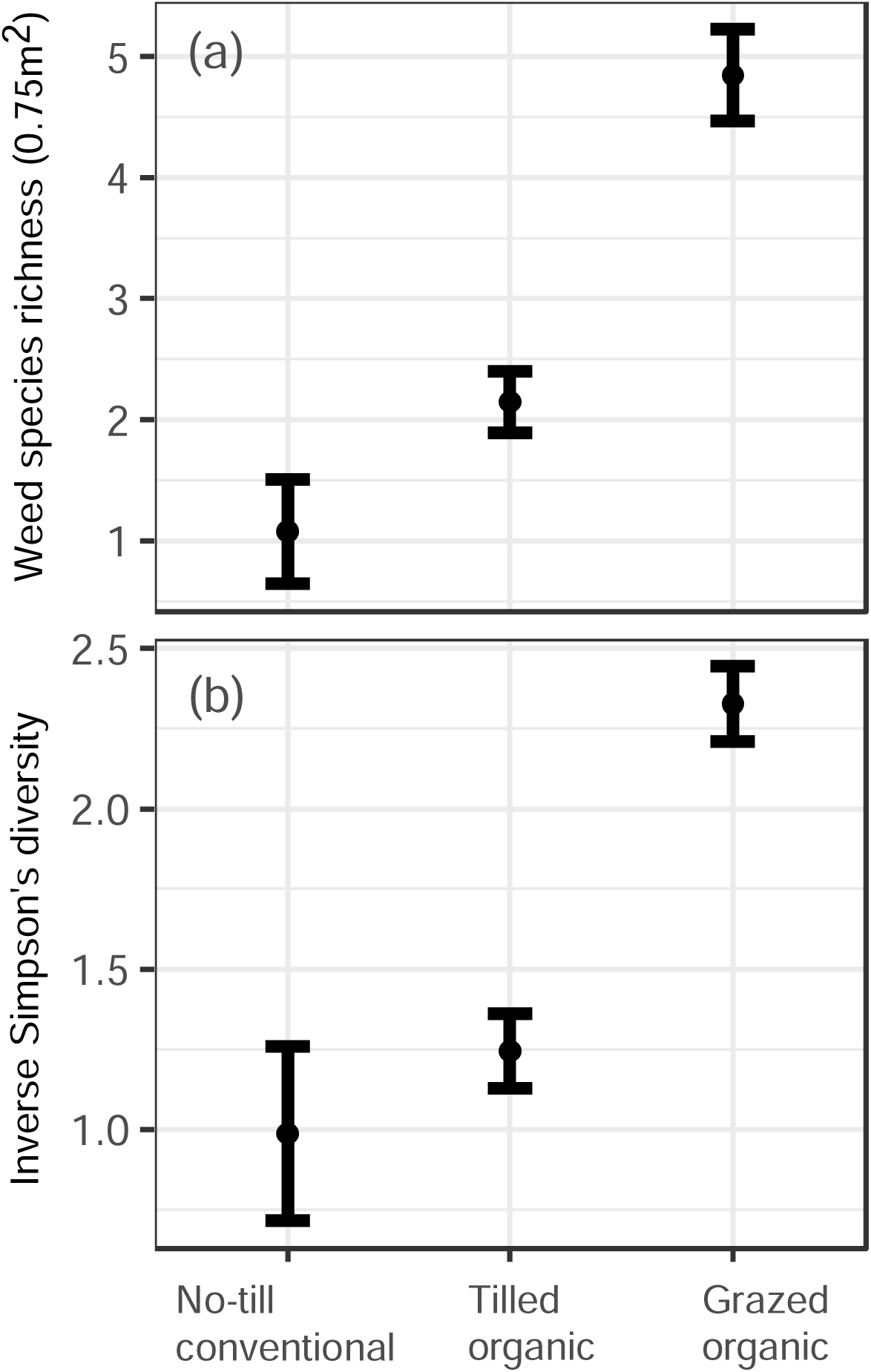
(a) Mean weed species richness, and (b) mean inverse Simpson’s diversity in 0.75 m^2^ split plots across three different cropping systems. Error bars indicate SE of the mean.

Weed biomass varied among cropping systems and years but was not affected by manipulated climate conditions (Table 1 & Fig 2). Overall across years, the lowest average weed biomass was sampled in the no-till conventional system (average <1 g of dry biomass ±0.02), followed by the tilled organic system (average 17 g of dry biomass ±10.3 SE), and the grazed organic system (average 38 g ±23.4 SE). Weed biomass was 10 times greater in 2017 than in 2016 in the tilled organic and grazed organic cropping systems (Fig 2). There were no differences in weed biomass among climate conditions within cropping systems, because of the large variation observed in the tilled organic and grazed organic systems. The exception was among ambient and warmer and drier conditions in 2016 in the tilled organic farming system (df=85, t=-2.36, P=0.05; Fig 2).

**Fig 2.**
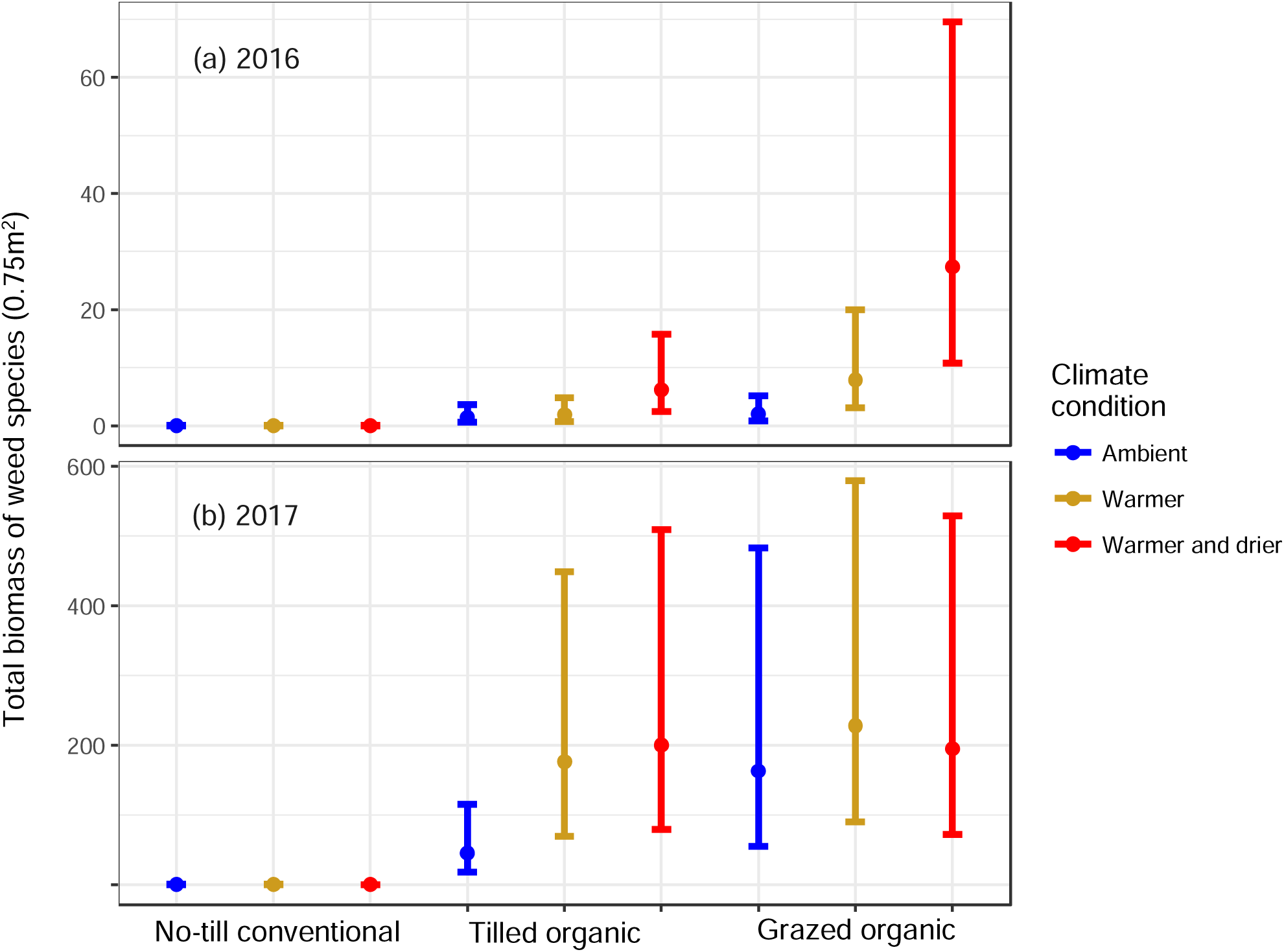
Estimated means of total biomass of weeds in 0.75 m^2^ split-plots in 2016 (a), and 2017 (b) by farming management system and climate condition. Error bars indicate SE of the mean.

**Table 1.** Type III Analysis of Variance of aboveground dry weed biomass in response cropping system and climate conditions, year of the experiment, and their interactions. Satterthwaite approximation was used to estimate degrees of freedom, and experimental block was fit as a random factor.

|  | Sum of<br>square<br>s | Mean<br>Square | Numerator<br>degree of<br>freedom | Denominat<br>or degrees<br>of freedom | F -<br>value | P<br>value |
| --- | --- | --- | --- | --- | --- | --- |
| Cropping system | 1021.6 | 510.79 | 2 | 85.02 | 139.0 | <0.001 |
| Climate condition | 5.47 | 2.74 | 2 | 85.02 | 0.7 | 0.478 |
| Year | 193.51 | 193.51 | 1 | 85.02 | 52.7 | <0.001 |
| Cropping system x Climate<br>condition | 29.66 | 7.41 | 4 | 85.02 | 2.0 | 0.099 |
| Cropping system x Year | 35.98 | 17.99 | 2 | 85.02 | 4.9 | 0.010 |
| Climate condition x Year | 4.68 | 2.34 | 2 | 85.02 | 0.6 | 0.532 |
| Cropping system x Climate<br>condition x Year | 7.31 | 1.83 | 4 | 85.02 | 0.5 | 0.738 |

The dissimilarity of weed species composition based on total weed biomass varied in response to cropping system and climate condition (Table 2) and was best explained by cropping system, which accounted for 16% of total variation in species composition. Climate condition accounted for 3% of total variation (Table 2). The tilled organic and grazed organic cropping systems were more variable in ordinal space, because of a greater diversity and variability in biomass (Fig. 2 and Fig 3). The no-till conventional system had the least variation in composition among the systems (see the ellipses in Fig 3) and was nested within one standard deviation of grazed organic system. Climate conditions did not indicate consistent shifts in weed communities within the cropping systems. The relative abundance of species differed across cropping systems. Specifically, *Thlaspi arvense* L dominated the weed communities sampled in the tilled organic system (Appendix Table S2). In contrast, in the grazed organic and no-till conventional system, *B. tectorum* was one of the most abundant species (Appendix Table S2).

**Fig 3.**
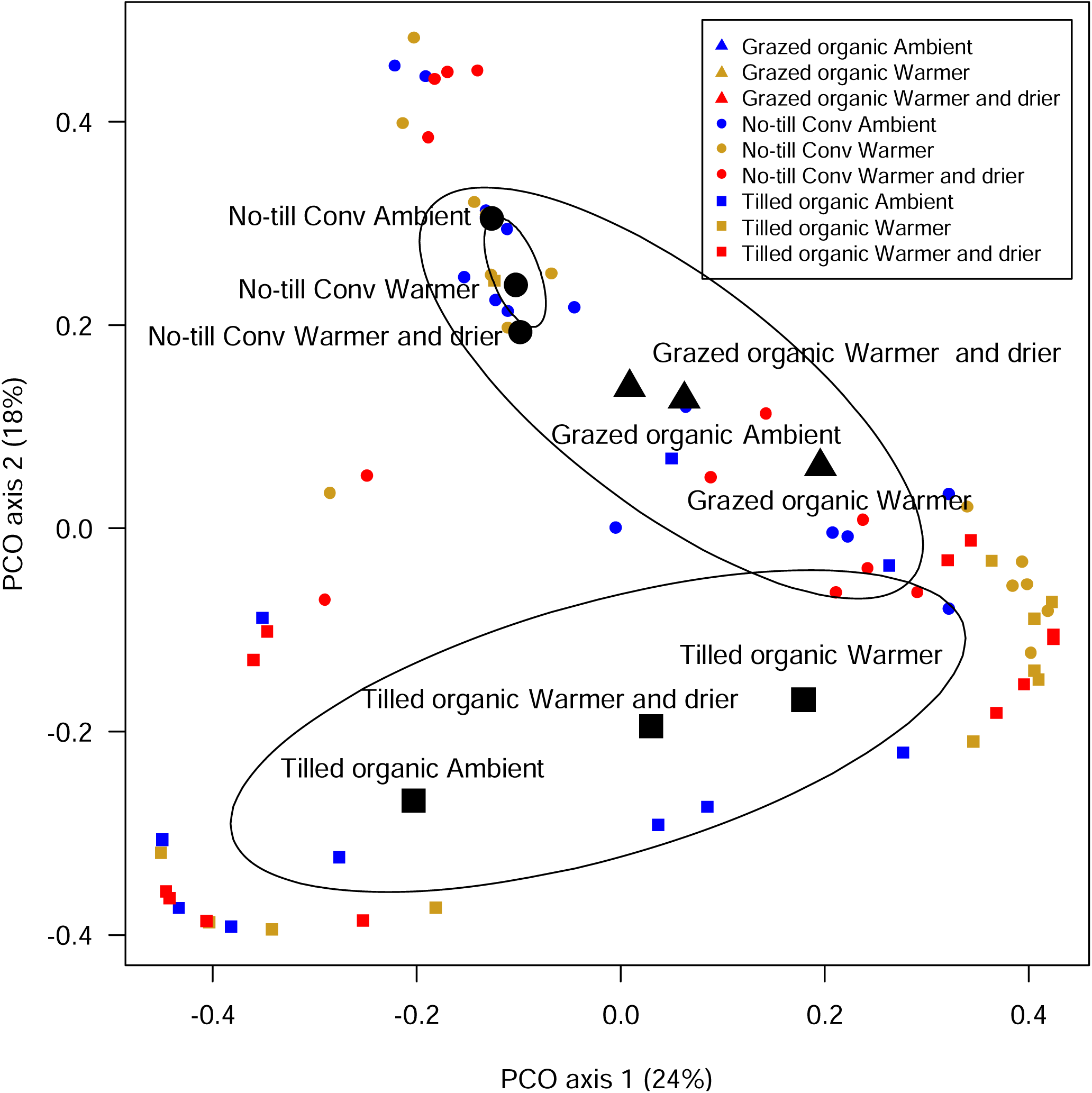
Principal coordinates ordination bi-plot of the dissimilarity of weed species biomass grouped by climate condition and cropping system. Ellipses indicate the standard deviation of centroids of cropping systems. Dissimilarity was calculated using the Bray-Curtis index based on biomass of weed species in each split-plot.

**Table 2.**
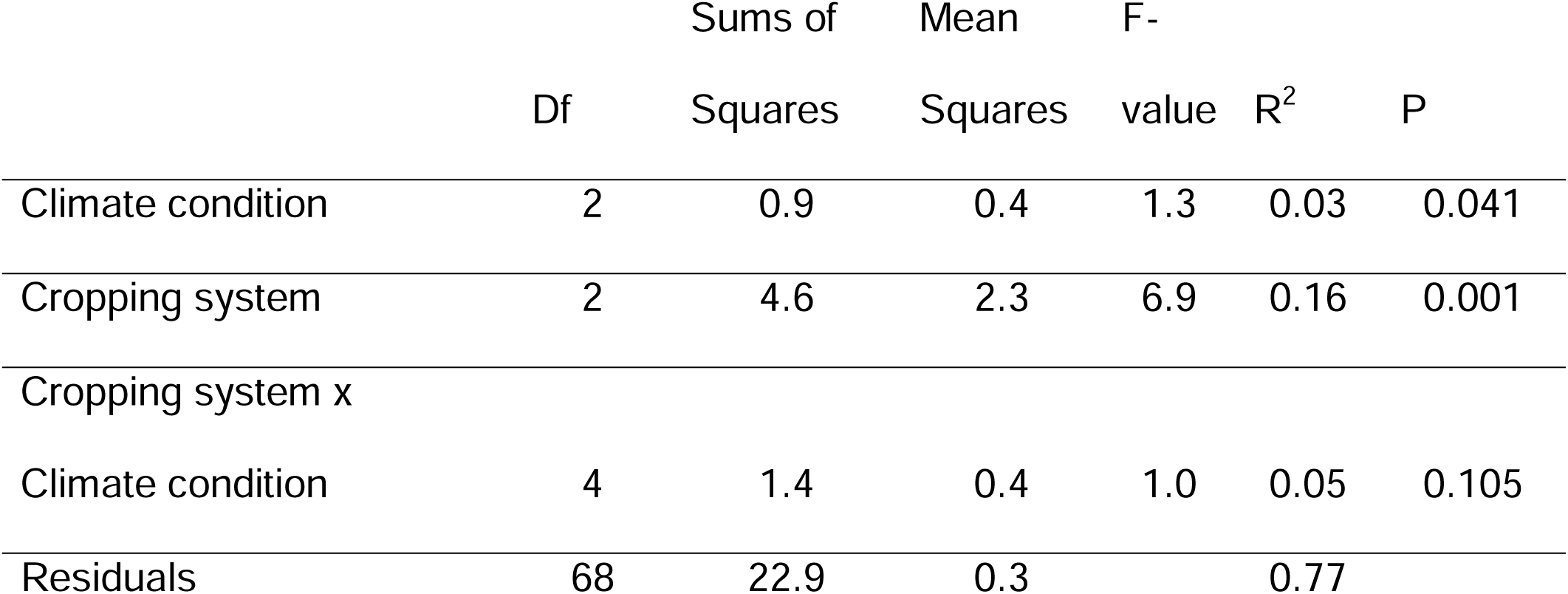
PERMANOVA of weed species composition in response to climate conditions and cropping system based on dissimilarity of species biomass using the Bray-Curtis index. Year was used to constrain variation.

### Wheat yield

Winter wheat yield varied among cropping systems, climate conditions, and years (Table 3). Winter wheat yield was lowest in the grazed organic system and highest in the no-till conventional farming system (Table 3 and 4, Fig. 4). Overall, across the three cropping systems yield of winter wheat declined by about 33% in response to warmer and drier conditions, when compared to the ambient climate conditions (Fig 3), but the response varied across cropping systems and years. Specifically, in the no-till conventional system, increased temperatures and drier conditions associated with a reduction in winter wheat yields in both years of the experiments (Table 4). In contrast, winter wheat yields in the grazed organic and tilled organic systems were reduced by warmer and drier conditions only in the second year of this study. (Table 4, & Fig. 4).

**Table 3.** Type III Analysis of Variance of wheat yield in response to cropping system, climate conditions, year of the experiment, and their interactions. Satterthwaite approximation was used to estimate degrees of freedom.

|  | Sum of<br>squares | Mean<br>Square | N df | D df | F -<br>value | P value |
| --- | --- | --- | --- | --- | --- | --- |
| Cropping system | 98.9 | 49.4 | 2 | 83.1 | 31.8 | <0.001 |
| Climate condition | 65.6 | 32.8 | 2 | 83.1 | 21.1 | <0.001 |
| Year | 26.9 | 26.9 | 1 | 83.1 | 17.3 | <0.001 |
| Cropping system x Climate<br>condition | 10.1 | 2.5 | 4 | 83.1 | 1.6 | 0.177 |
| Cropping system x Year | 31.0 | 15.5 | 2 | 83.0 | 10.0 | <0.001 |
| Climate condition x Year | 40.2 | 20.1 | 2 | 83.1 | 12.9 | <0.001 |
| Cropping system x Climate<br>condition x Year | 12.0 | 3.0 | 4 | 83.1 | 1.9 | 0.115 |

**Table 4.** Post-hoc Tukey test comparisons of wheat yield across climate conditions within cropping systems

| **2016** |  |  |  |  |  |
| --- | --- | --- | --- | --- | --- |
| **No-till conventional** | contrast estimate | SE | df | t value | P value |
| Ambient - Warmer | 0.86 | 0.72 | 83 | 1.2 | 0.46 |
| Ambient- Warmer and drier | 1.56 | 0.72 | 83 | 2.2 | 0.08 |
| Warmer - Warmer and drier | 0.70 | 0.72 | 83 | 1.0 | 0.60 |
| **Tilled organic** |  |  |  |  |  |
| Ambient - Warmer | -0.01 | 0.72 | 83 | -0.01 | 0.99 |
| Ambient- Warmer and drier | 0.23 | 0.72 | 83 | 0.33 | 0.94 |
| Warmer - Warmer and drier | 0.24 | 0.72 | 83 | 0.34 | 0.94 |
| **Grazed organic** |  |  |  |  |  |
| Ambient - Warmer | -1.49 | 0.81 | 84 | -1.8 | 0.16 |
| Ambient- Warmer and drier | -0.16 | 0.81 | 84 | -0.2 | 0.98 |
| Warmer - Warmer and drier | 1.33 | 0.72 | 83 | 1.8 | 0.16 |
| **2017** |  |  |  |  |  |
| **No-till conventional** |  |  |  |  |  |
| Ambient - Warmer | 1.32 | 0.72 | 83 | 1.8 | 0.16 |
| Ambient- Warmer and drier | 2.85 | 0.76 | 83 | 3.8 | 0.001 |
| Warmer - Warmer and drier | 1.52 | 0.76 | 83 | 2.0 | 0.12 |
| **Tilled organic** |  |  |  |  |  |
| Ambient - Warmer | 3.82 | 0.72 | 83 | 5.3 | <.0001 |
| Ambient- Warmer and drier | 4.73 | 0.72 | 83 | 6.6 | <.0001 |
| Warmer - Warmer and drier | 0.90 | 0.72 | 83 | 1.3 | 0.42 |
| **Grazed organic** |  |  |  |  |  |
| Ambient - Warmer | 1.60 | 0.76 | 83 | 2.1 | 0.09 |
| Ambient- Warmer and drier | 2.67 | 0.76 | 83 | 3.5 | 0.002 |
| Warmer - Warmer and drier | 1.07 | 0.79 | 83 | 1.4 | 0.37 |

**Fig 4.**
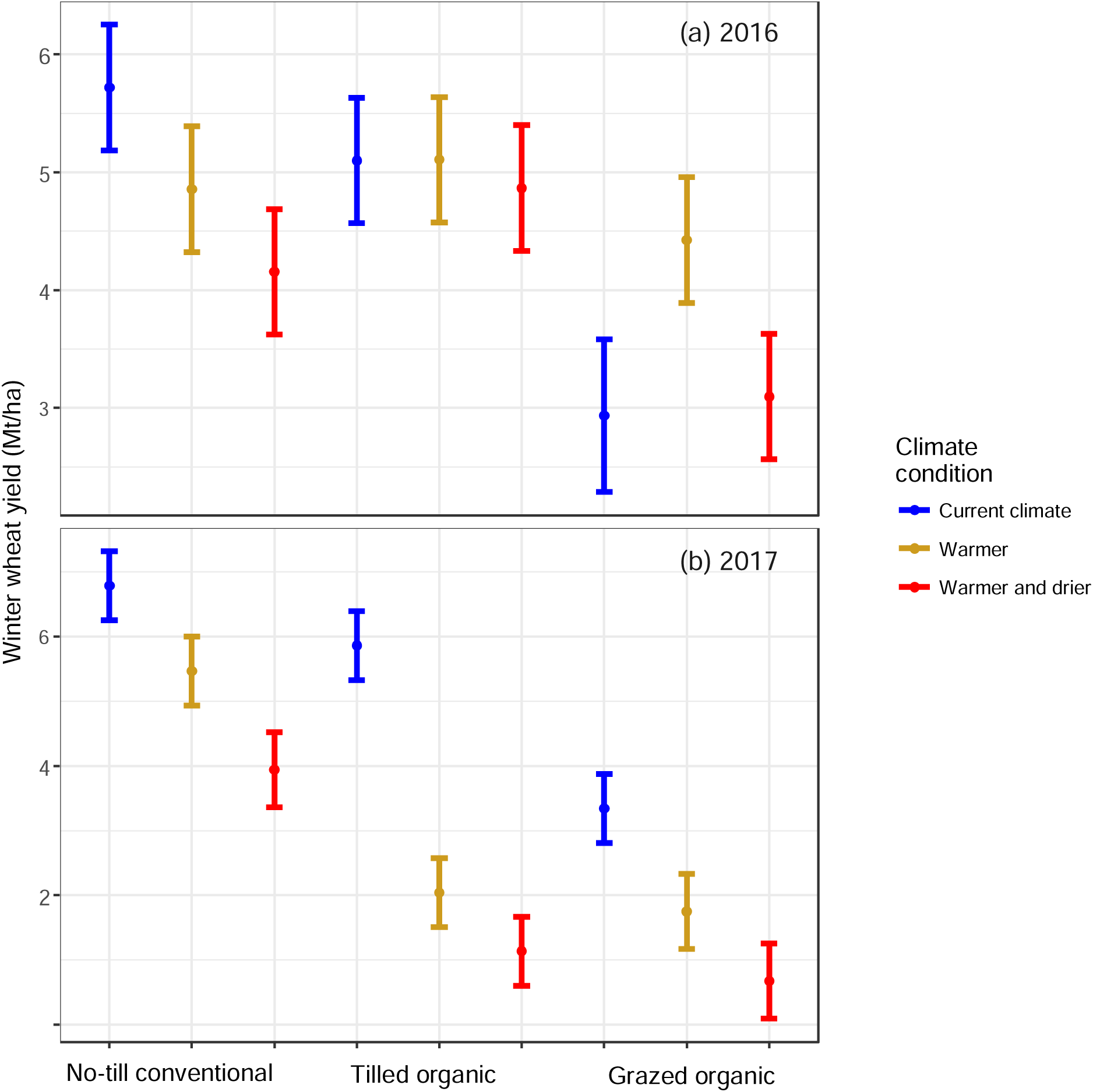
Wheat yield declined in response to warmer and drier conditions and varied by farming management systems and among years.

## Discussion

Population growth, coupled with shifts in consumer and market demands make it fundamental to understand how cropping systems respond to biological stressors and unprecedented climatic conditions (Searchinger et al., 2019). In this paper, we assessed how predicted warmer and drier climate conditions could impact weed communities and crop yields across three contrasting small grain cropping systems. This knowledge is essential as up to 60% of the global wheat production could be impacted by predicted warmer and drier climates (Trnka et al., 2019). Additionally, there is an urgent need to develop cropping systems that reduce their reliance on external inputs to manage pests and secure yields (Snapp et al., 2015, Reganold & Wachter, 2016, Peterson et al., 2018). Our results confirm previous studies that cropping systems have an overriding impact on weed communities’ characteristics and associated crop yields (Pollnac et al., 2009, Menalled et al., 2016, Ball et al., 2019). Also, also similar to previous studies (Lanning et al., 2010, Asseng et al., 2015, Morgounov et al., 2018), winter wheat yield declined in response to warmer and drier conditions, but the response varied across cropping systems. In the conventional no-till system, where there was almost no weed biomass, we observed a consistent decline in winter wheat yields in response to warmer and drier conditions regardless of the almost absent crop-weed competition. However, yield responses to warmer and drier conditions were more variable in the weedier tilled organic and grazed organic systems, supporting previous observations that the mechanisms driving crop-weed competition vary across cropping systems (Ryan et al., 2009, Smith et al., 2010, Storkey & Neve, 2018).

### Weed communities across cropping systems and climate conditions

Weed biomass and community composition varied across the three cropping systems in our study, which similar to findings that assessed changes in weed communities in conventional and organic cropping systems (Davis et al., 2005), and among different organic systems (Ball et al., 2019). Specifically, more weed biomass and increased species richness and diversity were observed in the two organic systems compared with the no-till conventional cropping system and the tilled organic system had less weed diversity biomass compared to the grazed organic system. The perennial weeds *Cirsium arvense* and *Taraxacum officinale* were more abundant in the grazed organic compared with the other cropping systems. Perennial weed species are a particular management challenge in organic cropping systems (Baker & Mohler, 2015, Orloff et al., 2018). While integrated weed management practices appears to be the most promising approach to manage perennial weeds in organic cropping systems (Orloff et al., 2018), our results highlight the difficulty of reducing tillage intensity in in the absence of synthetic herbicides.

While previous comparisons of weed communities between organic and conventional systems indicated clear shifts in species composition across cropping system (Menalled et al., 2001, Davis et al., 2005), we observed similar species composition between the reduced till grazed organic system and no-till conventional cropping systems. The no-till conventional community had lower diversity and biomass and was a subset of the grazed organic community. These weed communities were different from those sampled in tilled organic cropping system. In these two systems weed communities were dominated by *Bromus tectorum*, a species that is known to flourish in reduced tillage cereal cropping systems (Young et al., 2014). *Thlaspi arvense*, a winter annual species commonly found in organic small grain cropping systems of the Northern Great Plains (Adhikari & Menalled, 2018) was the most common weed species sampled in the tilled organic system, and was absent in the no-till conventional and a minor component of the grazed organic system. For 36 months prior to this study, tillage use was eliminated in the grazed organic and no-till conventional systems, underscoring the important effect tillage has on the assembly weed communities (Smith & Gross, 2007).

We detected small changes in weed communities in response to the climate conditions manipulation, probably due to the relatively short-term nature of the study, and that the relative change in soil temperature and moisture were with range of climate conditions experienced by wheat in the Northern Great Plains, a short coming of the in situ climate manipulations (Kreyling et al., 2017). We detected small trends to increasing biomass in the warmer and drier conditions in the organic systems, but this result was partially masked by the large split-plot to split-plot l variation caused by specific weed species such as *Lactuca serriola* and *Sisymbrium altissimum*.

### Climate conditions, cropping systems, and wheat yields

In arid and semi-arid environments, increased growing season stress and reduced kernel quality associated with climate change has and will continue to reduce wheat yields (Asseng et al., 2011, Asseng et al., 2015), consistent with our findings. Research on the effects of climate change has focused on stress associated with yield loss (Schlenker & Roberts, 2009), but recent research has focused on the use of agroecological management to design climate-resilient cropping systems (Altieri et al., 2015, Peterson et al., 2018). Organic systems have been hypothesized to be more resilient to climate change, yet plant-soil feedbacks and weed biomass in this study varied among organic systems indicating resilience varies among different organic cropping systems (Scialabba & Müller-Lindenlauf, 2010, Seipel et al., 2019).

In the studied organic systems, wheat yields did not decline in warmer and drier conditions during the first year of the study, but wheat yield were reduced in the second year. In contrast, though yields were highest in the no-till conventional system, they were consistently reduced in the manipulated warmer and drier conditions. The reason for the differences in yield decline in response to climate and weed biomass crop weed competition could be because of differences crop-weed competition observed between organic and conventional systems (Ryan et al., 2009, Stratonovitch et al., 2012), and could reflect different biotic interactions among wheat and soil microbial communities (Seipel et al., 2019, Ishaq et al., 2020).

### Conclusion

In this study, we demonstrated that weeds, cropping systems, and climate conditions interact to affect weed communities and winter wheat yields in the semi-arid Northern Great Plains. While studies have assessed how abiotic conditions will affect weed communities and wheat yields, this study shows that cropping systems have a direct impact and interact with climate conditions. Climate change will complicate weed management strategies but cropping system management can be used to design resilient cropping systems.

## Acknowledgements

This study was funded by a grant from USDA-ORG grant 2015-51106-23970 and USDA-Agriculture and Food Research Initiative - Foundational and Applied Science Program grant 1014774. We would like to thank Perry Miller, Jeff Holmes, and Devon Ragen for their help in managing the research site.

## Supporting Information

Additional Supporting Information may be found in the online version of this article:

Table S1. Monthly total precipitation and mean temperature during the field experiment

Figure S1. Images of split-plots in in the field experiment

Figure S2. Mean daily soil temperature 5 cm below the surface

Figure S3. Variation in soil moisture under ambient, warmer, and warmer and drier climate conditions

Table S2. The relative abundance of weeds species recorded

