## Supplemental information for "Weed communities and wheat yield are modified by cropping systems and climate conditions"

**Supplemental information Seipel et al.**

Table S1. Average monthly temperature and total monthly precipitation from PRISM[1] during important winter wheat growth stages in 2016 and 2017 when the experiment was conducted and the 30-year average from 1981-2010.

| Month | 2016 | | 2017 | | 1981-2010 Average | |
|---|---|---|---|---|---|---|
| | Average Temp (°C) | Total Precip (mm) | Average Temp (°C) | Total Precip (mm) | Average Temp (°C) | Total Precip (mm) |
| March | 3.7 | 48 | 5.1 | 42 | 1.6 | 34 |
| April | 7.9 | 53 | 6 | 93 | 5.8 | 57 |
| May | 9.8 | 68 | 10.6 | 69 | 10.4 | 79 |
| June | 16 | 28 | 15.2 | 69 | 14.5 | 79 |
| 4-month | 9.4 | 197 | 9.2 | 273 | 8.1 | 249 |

[1]PRISM Climate Group. (2018). Oregon State University. *http://prism.oregonstate.edu*.

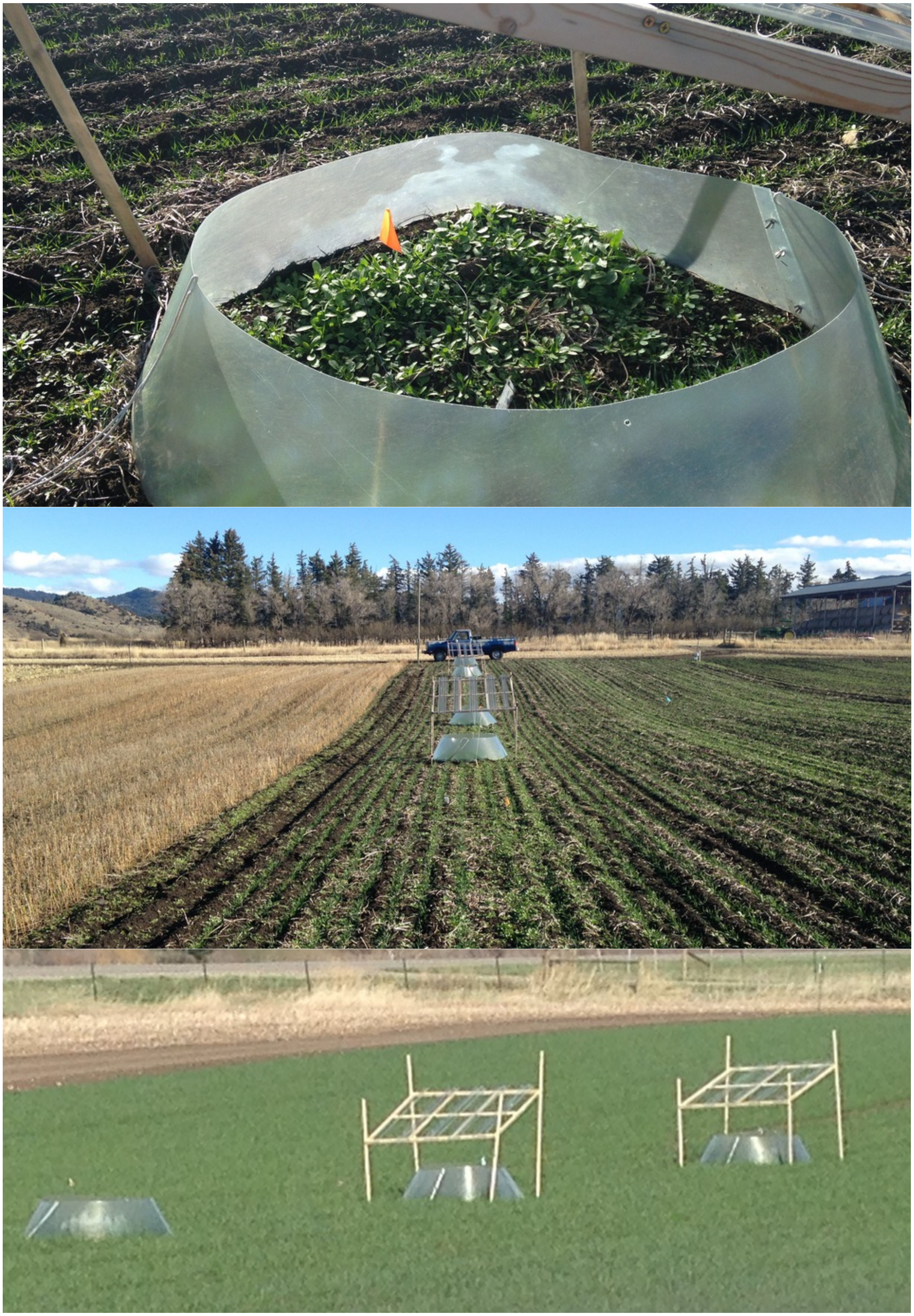

Figure S1. Photos of the experiment in the field with climate manipulation structures.

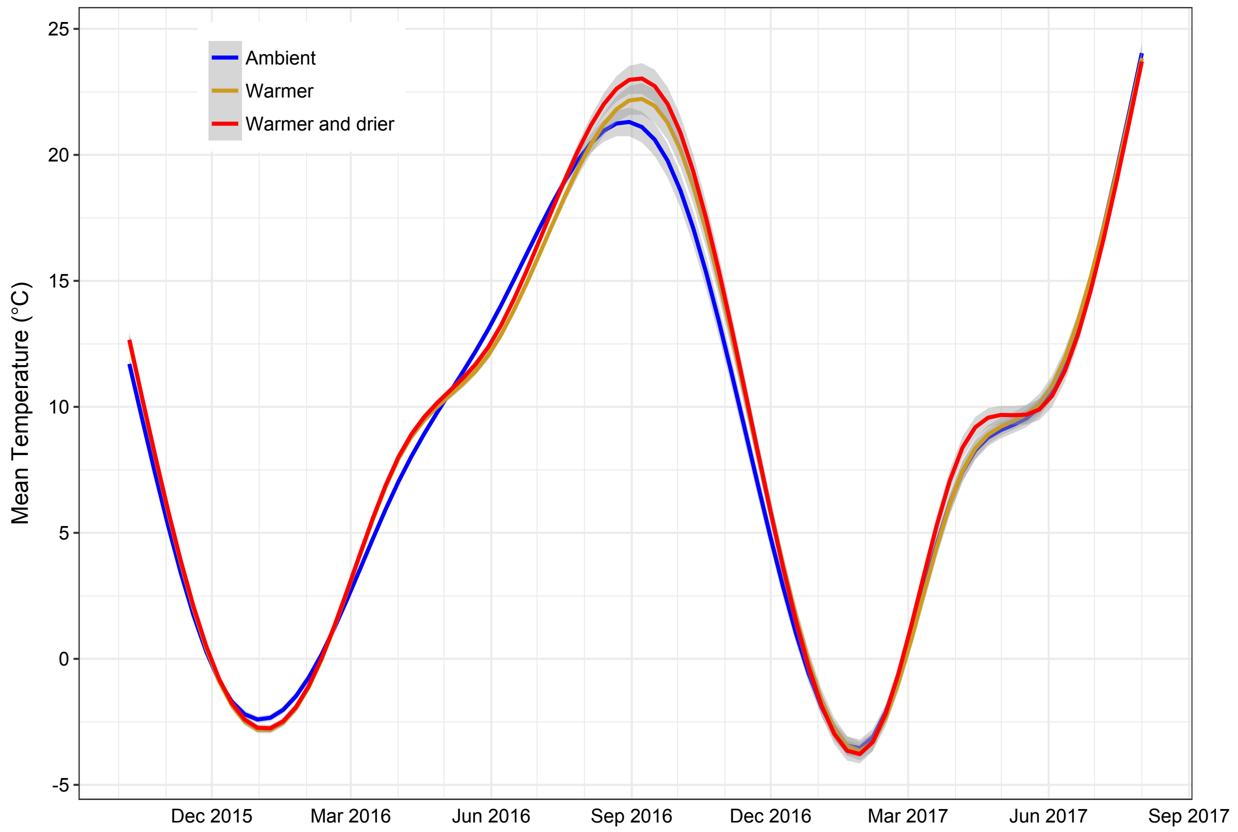


Figure S2. Mean daily soil temperature 5 cm below the surface in winter wheat fields with ambient conditions and climate manipulations that made the soil temperature warmer using open-top chambers, and warmer and drier using open-top chambers and rainout shelters that block 50% of the area above the plots.

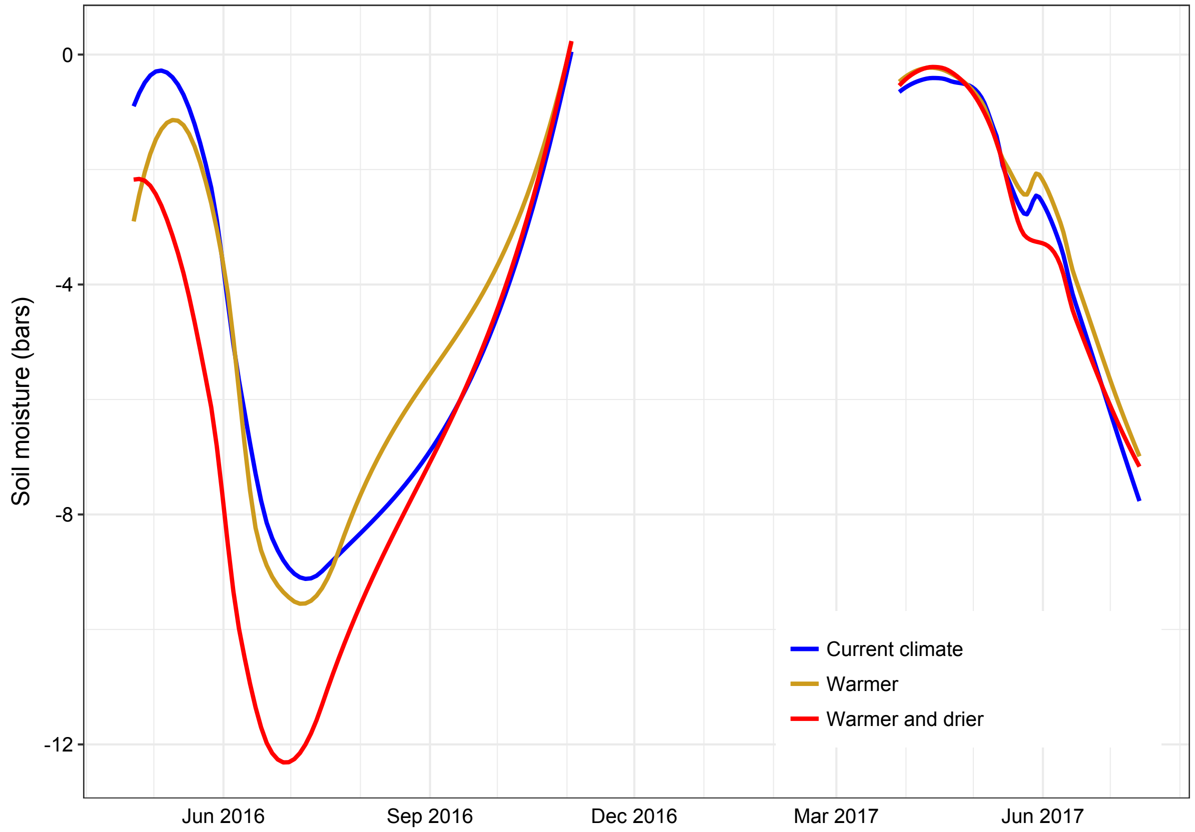


Figure S3. Variation in soil moisture under ambient, warmer, and warmer and drier climate conditions.

Table S2. The relative abundance of weeds species recorded in 26 $m^2$ in each cropping system.

| No-till conventional | | Grazed organic | | Tilled organic | |
|---|---|---|---|---|---|
| Weed species | total dry biomass (g) | Weed species | total dry biomass (g) | Weed species | total dry biomass (g) |
| *Bromus tectorum* | 17.83 | *Bromus tectorum* | 1429.8 | *Thlaspi arvense* | 2221.3 |
| *Lactuca serriola* | 16.86 | *Sisymbrium altissimum* | 1296.0 | *Lactuca serriola* | 615.3 |
| *Chenopodium album* | 0.02 | *Thlaspi arvense* | 1092.0 | *Asperugo procumbens* | 87.7 |
| | | *Lactuca serriola* | 622.7 | *Capsella bursa-pastoris* | 27.2 |
| | | *Asperugo procumbens* | 476.1 | *Cirsium arvense* | 22.4 |
| | | *Taraxacum officinale* | 196.6 | *Camelina microcarpa* | 10.3 |
| | | *Capsella bursa-pastoris* | 150.7 | *Sisymbrium altissimum* | 7.4 |
| | | *Tragopogon dubius* | 137.5 | *Tragopogon dubius* | 3.5 |
| | | *Melilotus officinalis* | 89.0 | *Chenopodium album* | 1.1 |
| | | *Cirsium arvense* | 71.0 | *Taraxacum officinale* | 0.9 |
| | | *Trifolium pratense* | 32.9 | *Bromus tectorum* | 0.6 |
| | | *Bromus japonicus* | 15.8 | | |
| | | *Xanthium strumarium* | 7.2 | | |
| | | *Dactylis glomerata* | 7.0 | | |
| | | *Descurainia sophia* | 6.5 | | |
| | | *Agropyron cristatum* | 4.7 | | |
| | | *Galium aparine* | 3.8 | | |
| | | *Lotus corniculatus* | 3.0 | | |
| | | *Medicago sativa* | 0.6 | | |
| | | *Avena fatua* | 0.5 | | |
